# The protection of Gα_z_-null NOD mice from hyperglycemia is sexually dimorphic and only partially β-cell autonomous

**DOI:** 10.1101/2021.02.08.430304

**Authors:** Rachel J. Fenske, Darby C. Peter, Haley N. Wienkes, Michael D. Schaid, Austin Reuter, Kathryn A. Carbajal, Michelle E. Kimple

## Abstract

The mechanisms that underlie the β-cell pathophysiology of Type 1 Diabetes (T1D) are not fully understood. Our group has defined the unique heterotrimeric G protein alpha-subunit, Gα_z_, as a key negative regulator of β-cell signal transduction pathways. Non-obese diabetic (NOD) mice lacking Gα_z_ throughout the body are protected from developing T1D-like hyperglycemia. To determine whether this phenotype is β-cell autonomous, we generated and validated a β-cell-specific Gα_z_ knockout (βKO) on the NOD background and characterized the phenotype of female and male cohorts. Long-term hyperglycemia incidence was lower in Gα_z_ βKO mice as compared to wild-type (WT) controls, but, unlike global Gα_z_ knockout mice, this protection was incomplete. While young male and female Gα_z_ βKO NOD mice had improved glucose tolerance, WT NOD males were significantly less glucose tolerant than females, and only female Gα_z_ βKO mice retained improved glucose tolerance at 28-29 weeks of age. Conversely, β-cell-specific Gα_z_ loss only influenced insulitis in 28-29-week old male NOD mice, a phenotype correlating directly with body burden of glucose during oral glucose challenge. Using surrogates for β-cell function and apoptosis, the partial penetrance of euglycemia in Gα_z_ βKO NOD was best explained by an early failure to up-regulate β-cell proliferation. We conclude β-cell Gα_z_ is an important regulator of the sexually-dimorphic T1D-like phenotype of NOD mice. Yet, other factors must be important in imparting full protection from the disease.

## Introduction

Type 1 diabetes (T1D) is an autoimmune disease that is growing in prevalence in both youth and adults globally. Although T1D is characterized primarily as a loss of β-cell mass as a direct result of immune-mediated β-cell destruction, a growing body of evidence suggests the β-cell is not a passive player in this dysfunctional immune response. Β-cell factors, both secreted and intra-cellular signaling molecules, are thought to be pathological contributors to disease onset and progression. The interaction of β-cells with other islet cell types and immune cells in their microenvironment can drive or inhibit insulin secretion. Β-cells secrete autoantigens, primarily preproinsulin, that are recognized by cytotoxic T lymphocytes (CTL) and trigger targeting of the β-cell for attack (1,2). Β-cells are also prone to endoplasmic reticulum (ER) stress due to their high secretory demand, which amplifies the immunogenicity of autoantigen secretion (3). Β-cells must make and store high quantities of insulin in order to dynamically regulate insulin secretion in response to glucose. In order to maintain high secretory capacity, β-cells must be able to upregulate pathways that promote insulin secretion.

Cyclic adenosine monophosphate (cAMP) is a central amplifier of β-cell insulin secretion (4). Cyclic AMP levels are maintained through a balance of cAMP production by adenylyl cyclases (AC), and cAMP degradation by phosphodiesterases (PDE) (5,6). Type 2 diabetes therapeutics that work, in part, by stimulating β-cell cAMP production (i.e., glucagon-like peptide 1 receptor agonists: GLP1-RAs) have been proven clinically effective in improving β-cell function, thereby lowering HbA1c (7,8). With evidence primarily from rodent models for stimulating β-cell proliferation and survival, their usefulness as adjuvant therapies in patients with T1D is currently being investigated (9–12).

We have previously established the alpha-subunit of the heterotrimeric G_z_ protein, Gα_z_, as an important negative regulator of β-cell cAMP production and downstream function, proliferation and/or survival(13–17). Mice globally lacking Gα_z_ have enhanced glucose clearance and are protected from developing hyperglycemia and glucose intolerance in a number of mouse models of diabetes, including the Non-obese diabetic (NOD) model of spontaneous T1D (13–16). When co-administered a sub-therapeutic dose of the GLP1-RA, exendin-4, full-body Gα_z_ knockout mice (FΒKO) in the C57BL/6N background are fully protected from developing hyperglycemia after multiple low dose streptozotocin (MLD-STZ) induction of T1D (13). In both the NOD and MLD-STZ models, global loss of Gα_z_ is associated with improved β-cell function, reduced β-cell apoptosis, and increased β-cell replication (13,14). Furthermore, global Gα_z_ loss has no direct effect on NOD immune phenotype (14). As β-cell-specific Gα_z_ knockout mice (Gα_z_ βKOs) in the C57BL/6J background are equally as protected from MLD-STZ as FΒKO (13,14), the T1D protection phenotype appears best explained by a β-cell-centric model.

To confirm β-cell Gα_z_ as the driver of T1D-like pathophysiology in the NOD background, we back-crossed the Gα_z_ βKO mutation into the NOD background for 10 generations, confirming preservation of NOD loci. We next characterized disease progression and penetrance in both males and females, correlating this with different *in vivo* and *ex vivo* measurements of glucose control, β-cell function, and insulitis. Our results reveal an unexpected sex-specific effect of β-cell Gα_z_ loss on various aspects of the pathophysiology of the NOD phenotype. Furthermore, the incomplete protection of Gα_z_ βKO females from hyperglycemia is associated with a failure to up-regulate the β-cell proliferation, suggesting β-cell non-autonomous mechanisms are at play in the FΒKO.

## Materials and Methods

### Animals

The Gα_z_ βKO NOD strain was generated by crossing NOD mice with Gα_z_ βKO mice in the C57BL/6J strain, in which Cre is under the control of the rat insulin promoter (“InsPr-Cre” in (18)). The β-cell specificity of Gα_z_ loss was validated previously(14). NOD mice were backcrossed for more than 10 generations and SNP genotyped by the Jackson Laboratory to ensure full preservation of NOD loci. Breeding colonies were housed in a limited access, pathogen-free facility where all cages, enrichment, and water are sterilized prior to use on a 12 h light/dark cycle with ad libitum access to water and irradiated breeder chow (Teklad 2919, Envigo). Doble heterozygous crosses were performed to obtain Gα_z_ βKO and wild-type Cre-positive controls (WT). Experimental animals were identified by genotyping for the floxed *Gnaz* gene and Cre transgene as described previously (14). Gα_z_ FΒKO mice in the C57BL/6N background have been extensively characterized by our group (13,16,19) and were used in MLD-STZ experiments.

Blood glucose measurements were collected weekly from 4 weeks of age up to 35 weeks of age using a blood glucose meter (AlphaTRAK) and rat/mouse-specific test strips as previously described (14). At the end of the study, pancreases were collected for paraffin embedding using a well-established protocol (20). A separate cohort of mice was sacrificed at the 12-13 week time point and pancreases or islets were collected from females and males, respectively.

### Oral glucose tolerance tests (OGTTs)

Animals were fasted for 4-6 hours prior to glucose tolerance testing at both 12-13 weeks of age and 28-29 weeks of age for oral glucose tolerance tests (OGTTs). Glucose was given using an oral gavage at a dose of 2 g/kg BW. Blood glucose measurements were taken immediately prior to gavage and 5, 15, 30, 60, and 120 minutes following gavage. At 12-13 weeks of age, blood from male and female animals was collected using a lateral tail nick at baseline and 5 minutes after gavage. At 28-29 weeks of age, blood was only collected from female animals at baseline and 5 minutes post-gavage. Blood was spun down at 3000 g for 10 minutes at 4 degrees. Plasma was collected and insulin was measured using an Ultra Sensitive Mouse Insulin ELISA (Crystal Chem; catalog #90080) per the manufacturer’s instructions.

### Whole Pancreas Collection and Staining

Following dissection, pancreata were kept in 10% formalin for 48 hours before being transferred to 70% ethanol until paraffin fixation by the University of Wisconsin – Carbone Cancer Center Experimental Pathology Laboratory (University of Wisconsin Carbone Cancer Center Support Grant P30 CA014520). 10-micron serial sections were used in downstream analyses. Immune infiltration of the islet was determined using standard eosin (Polysciences Cat #17269) and hematoxylin (Sigma, GHS280) staining. Following staining, quantification of immune infiltration was accomplished using a numerical scoring system, as previously described (14). For insulin IHC to calculate β-cell fractional area, a 1:500 dilution of guinea pig anti-insulin primary antibody (Dako, discontinued) and 1:2000 goat anti-guinea pig HRP-coupled secondary antibody (Cell Signaling) were used, and slides processed and imaged as previously described (13,14,19). To determine Ki67-positive, insulin-positive cells, rabbit anti-Ki67 (Cell Signaling - 1:200) and guinea-pig anti-insulin (Abcam - 1:75) were applied to sections overnight at 4°C. After a PBS wash, slides were incubated for 30 min with 1:400 FITC-coupled donkey anti-rabbit IgG and 1:200 Cy3-coupled goat anti-guinea pig IgG (both from Jackson Immunoresearch). Slides were processed and imaged as previously described(19) In both imaging experiments, islets from the entire pancreas section area were quantified, with two sections quantified per mouse (N=3 mice/group).

### Low-dose STZ protocol for β-cell insult

10-12 week-old WT and Gα_z_ FΒKO NOD animals and 10-12 week-old WT Cre+ and Gα_z_ ΒKO NOD females were separated into groups to receive either STZ (50 mg/kg) or citrate buffer for 3 consecutive days, in order to induce a β-cell insult. STZ was made at a concentration of 7.5 mg/mL, protected from light, at least 2 hours prior to injection to give adequate equilibration time (21). Injection procedure was performed as previously describe, save for only 3 days vs. the standard 5 day MLD-STZ protocol (13).

### Statistical Analysis

Data are expressed as mean ± standard error of the mean unless otherwise noted. Data were compared by two-way analysis of variance or Student *t* test as appropriate and as described in the figure legends. A *P* value <0.05 was considered statistically significant. Statistical analyses were performed with GraphPad Prism version 8 (GraphPad Software, San Diego, CA).

## Results

### Loss of β-cell Gα_z_ is not sufficient to protect NOD mice from developing hyperglycemia

Random-fed blood glucose levels of WT and Gα_z_ βKO NOD mice were tracked from weaning (3-4 weeks of age) until 30 weeks of age (male) or 34 weeks of age (female). Hyperglycemia was classified as a random-fed blood glucose level of >250 mg/dl that remained elevated the following week. As has been well-characterized in the literature, the incidence of hyperglycemia was significantly lower in male NOD mice vs. female, with approximately 10% penetrance in males and 40% in females by study end (**Figure 1A,B**). Thirteen of 32 WT females had become hyperglycemic by 34 weeks of age compared to 4 of 14 βKO females, corresponding to a disease-free survival of 59.3% and 71.4%, respectively; a modest difference that was not statistically significant (**Figure 1A**). This is in contrast to the previously-published 100% disease-free survival of Gα_z_ FΒKO NOD females over the same timeframe, in the same animal facility(14). Two of 19 WT NOD males became hyperglycemic before 30 weeks of age, as compared to zero of 8 βKO males, with a disease-free survival of 89.4% and 100%, respectively. This effect size is similar to that observed with Gα_z_ ΒKO NOD males vs. WT (14); yet, with so few cases overall, it was not statistically-significant (**Figure 1B**)

**Figure 1:**
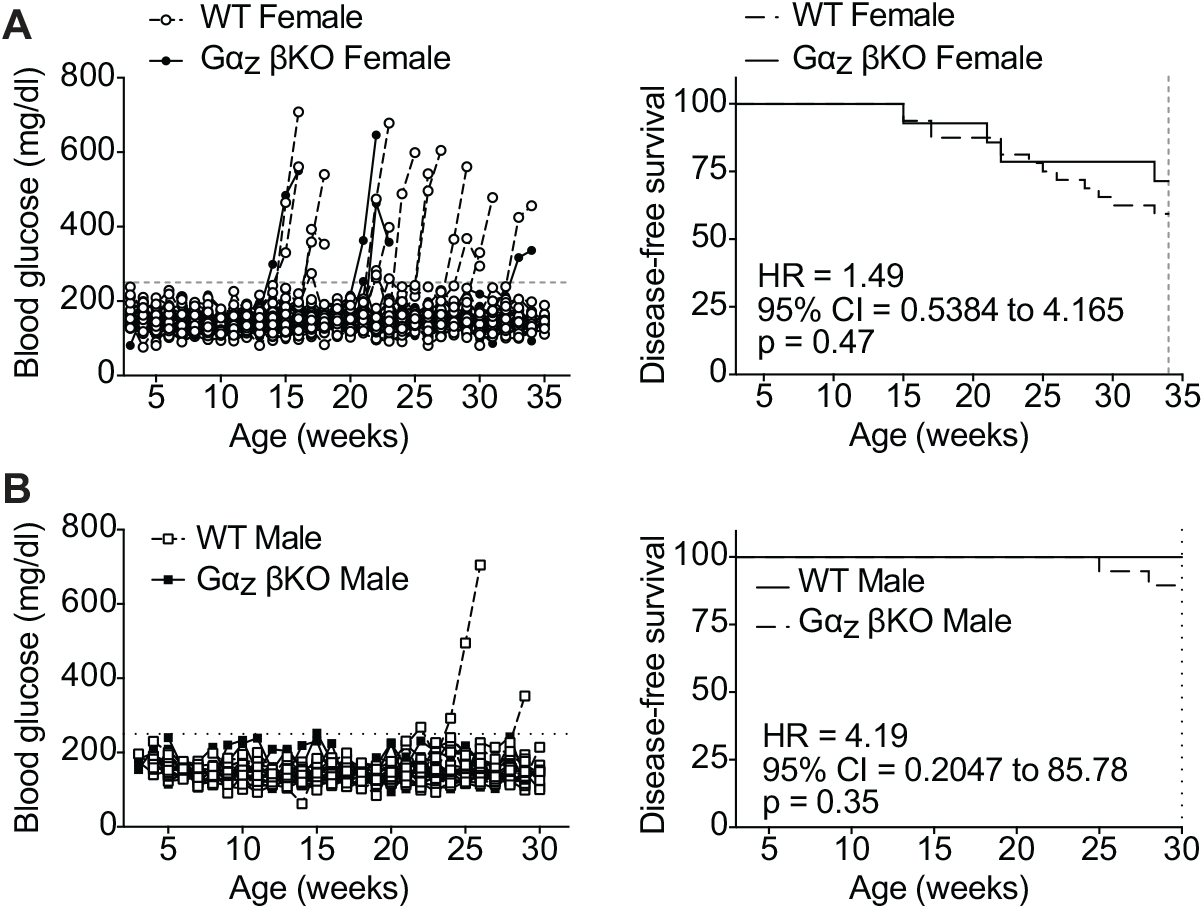
Β-cell loss of Gα_z_ only partially protected NOD mice from developing T1D-like hyperglycemia. A: Random fed blood glucose levels (left) and disease-free survival (right) of female WT and Gα_z_ βKO mice from weaning (3-4 weeks of age) until 34 weeks of age. B: Random-fed blood glucose levels (left) and disease-free survival (right) of male WT and Gα_z_ βKO mice from weaning (3-4 weeks of age) until 30 weeks of age. Disease-free survival was defined as random-fed blood glucose levels above 250 mg/dl for two consecutive weeks. Data in the right panels were compared via Log-rank (Mantel-Cox) test to determine whether the survival curves were significantly different (p < 0.05). Also shown are the log-rank hazard ratio and (HR) and 95% confidence interval (CI).

### Glucose tolerance is improved over the lifetime of NOD females when β-cell Gα_z_ is lost

Loss of glucose tolerance is one method by which to characterize emerging β-cell failure in T1D (22). OGTTs were performed at two time points over the course of the study: 12-13 weeks (before overt hyperglycemia in WT mice of either sex), and 28-29 weeks (full disease penetrance).

At 12-13 weeks of age, there was no statistically-significant difference in 4-6 h fasting blood glucose levels by sex or genotype (**Figure 2A**). Both male and female Gα_z_ βKO mice cleared glucose from their blood faster than WT controls, with the strongest effect at the 15 min time point (**Figure 2B,C**). Calculating the glucose area-under-the-curve (AUC) during OGTTs for showed a significant reduction by genotype in both sexes, as well as a significantly lower OGTT AUC in female NOD mice as compared to male, independent of genotype (**Figure 2D**). Plasma insulin measured at the 5 min time point during OGTTs showed a higher mean in βKO mice as compared to WT in both sexes, although none of means were significantly higher at 5 min than at zero (**Figure 2E**). Plotting the raw values for each mouse is also suggestive, although not confirmatory, of a secretion effect in Gα_z_ βKO mice (**Figure 2F**).

**Figure 2:**
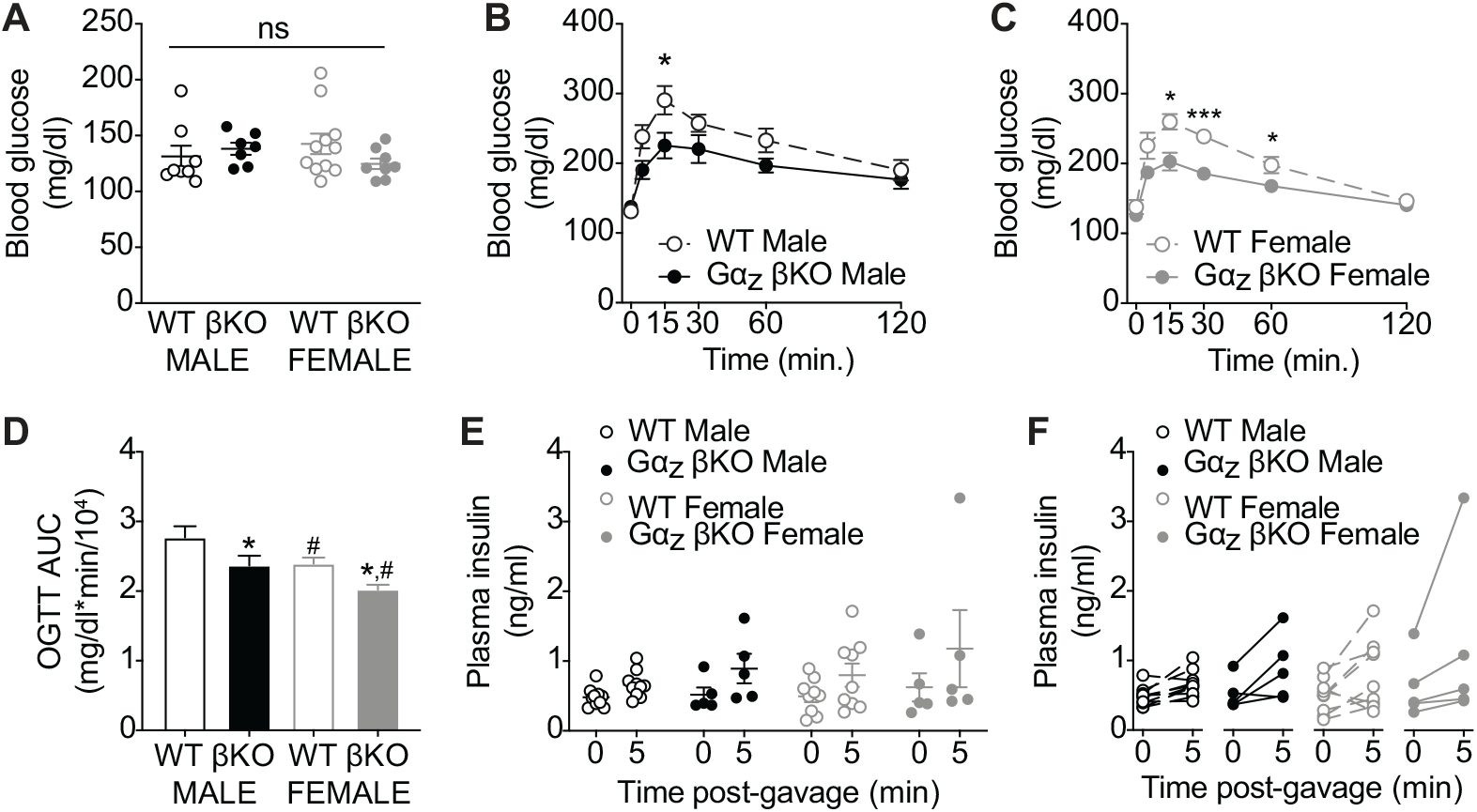
Prior to hyperglycemic onset, glucose tolerance is improved in Gα_z_ βKO male and female NOD mice. A: 4-6 fasting blood glucose levels of WT and βKO male and female NOD mice. N=7-11. B,C: Glucose excursions after oral challenge with 2 g/kg glucose in male (B) and female (C) NOD mice. N=7-11. D: Average area-under-the-curve of data shown in (B) and (C). E,F: Mean plasma insulin levels by group (E) and raw plasma insulin levels per mouse (F) during OGTTs. N=5-10. In all panels, data represent means ± SEM. In (A) and (D), data were compared by one-way ANOVA with Holm-Sidak test post-hoc to correct for multiple comparisons. In (B), (C), and (E), data were compared by two-way ANOVA with Holm-Sidak test post-hoc to correct for multiple comparisons. *p < 0.05, **p < 0.01, ***p < 0.001, and ****p < 0.0001 vs same-sex WT control; ^#^p < 0.05 male vs. female. If a comparison is not shown, it was not statistically-significant.

At 28-29 weeks of age, there were still no statistically-significant differences among the 4-6 h fasting blood glucose levels of male or female NOD mice, WT or βKO (**Figure 3A**). In contrast to the earlier age, only female βKO mice remained more glucose tolerant than their WT counterparts, with significant reductions in blood glucose levels 15 and 30 min after glucose challenge (**Figure 3B,C**). Interestingly, though, the biggest difference in male glucose tolerance at the 28-29 week vs. 12-13 week timepoint was a significant enhancement in glucose clearance in the WT cohort, as evidenced by comparison of OGTT AUCs (**Figure 3D, left**). This is in contrast to female WT NOD mice, who have almost identical OGTT AUCs at both young and older ages (**Figure 3D, right**). Comparing results between sexes, by 28-29 weeks of age WT male NOD mice are no longer less glucose tolerant than WT females, and there is no additive effect of the βKO (**Figure 3D**). Plasma insulin levels 5 min after glucose administration were not significantly different in male or female βKO mice as compared to WT, but, like 12-13 week data, there was no significant enhancement at 5 min vs baseline in any group (**Figure 3E**). Interestingly, plotting the raw data by individual mouse suggests a stronger plasma insulin release in 28-29-week old male vs. female NOD mice (**Figure 3F**): a trend not observed at 12-13 weeks of age.

**Figure 3:**
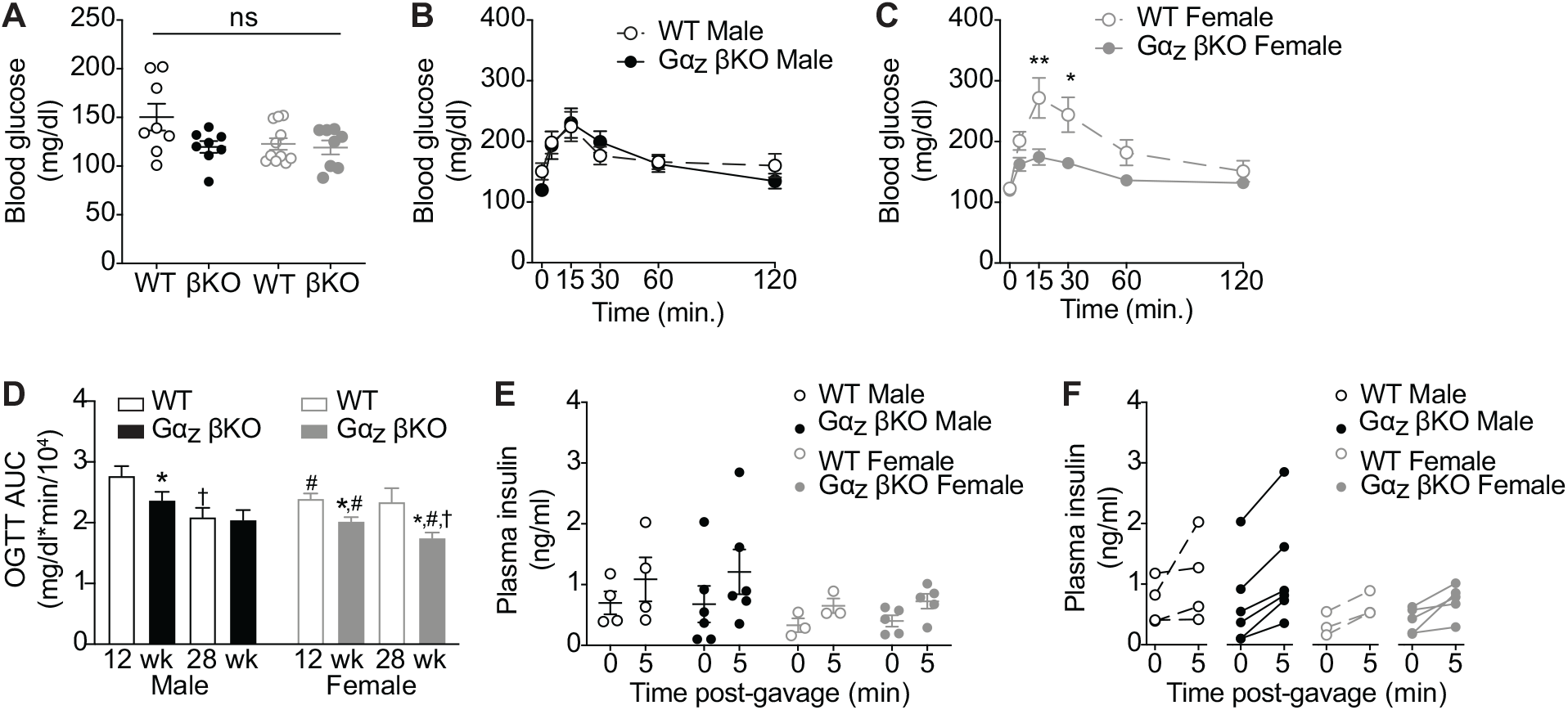
At 28-29 weeks of age, improved glucose tolerance of Gα_z_ βKO NOD mice is only maintained in females. A: 4-6 fasting blood glucose levels of WT and βKO male and female NOD mice. N=8-11. B,C: Glucose excursions after oral challenge with 2 g/kg glucose in male (B) and female (C) NOD mice. N-8-11. D: Average area-under-the-curve of data shown in (B) and (C), as compared to the AUC data shown for 12-13 weeks in Figure 2. N-7-11. E,F: Mean plasma insulin levels by group (E) and raw plasma insulin levels per mouse (F) during OGTTs. N=3-6. In all panels, data represent means ± SEM. In (A) and (D), data were compared by one-way ANOVA with Holm-Sidak test post-hoc to correct for multiple comparisons. In (B), (C), and (E), data were compared by two-way ANOVA with Holm-Sidak test post-hoc to correct for multiple comparisons. *p < 0.05 and **p < 0.01 vs. same sex, same age WT control; ^#^p < 0.05 male vs. female same-age control.^†^, p<0.05 for 28 wk. vs. 12 wk. within sex. If a comparison is not shown, it was not statistically-significant.

### Loss of β-cell Gα_z_ only improves insulitis in male NOD

Both male and female NOD mice display insulitis, which is a direct contributor to the β-cell dysfunction of T1D. Insulitis was quantified at study end (30 weeks – male and 34 weeks – female) by hematoxylin and eosin staining of pancreas slide sections according to a scoring system previously published(14). Interestingly, male Gα_z_ βKO mice had a significantly reduced insulitis than WT controls, whether represented as the mean islet immune cell infiltration score (**Figure 4A**) or the percentage of islets with each score (**Figure 4B**). There were no other significant differences by sex or genotype. For each mouse, its islet infiltration score was plotted against its 28-29 week OGTT AUC, and only in male mice did glucose tolerance correlate with insulitis (p = 0.021, r^2^ = 0.4637) (Figure 4D). To our knowledge, this sexually-dimorphic phenotype has not previously been described.

**Figure 4:**
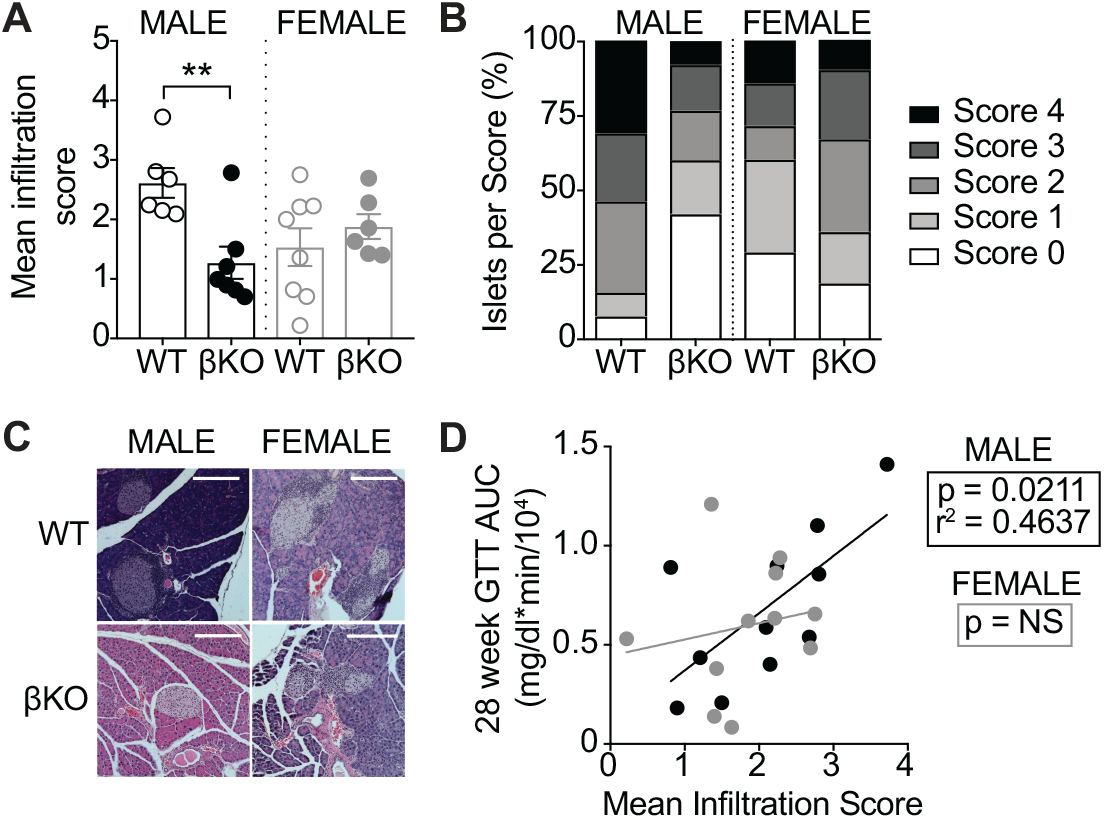
Loss of β-cell Gα_z_ only improves insulitis in male NOD. A,B: Severity of insulitis was quantified by (A) mean islet infiltration score per pancreas, or (B) as the percentage of islets with each score, both as described in (14). N=6-8. C: Representative islet images for the analysis shown in panels (A) and (B). Scale bar represents 200 microns. D: Mean infiltration score was plotted against 28-29 week-GTT AUC and a linear regression fit for males and females, exclusive of genotype. The goodness-of-fit (r^2^) and significance (p-value) for the linear regression are indicated. In (A), data were compared by one-way ANOVA with Holm-Sidak test post-hoc to correct for multiple comparisons. **p < 0.01 vs. WT.

### Β-cell loss of Gα_z_ did not alter β-cell area or proliferation

Β-cell fractional area was quantified at study end by insulin immunohistochemistry of pancreas slide sections counter-stained with hematoxylin. There were no significant differences in insulin-positive pancreas area by sex or by genotype (**Figure 1A**). This is in contrast to our previous findings of significantly elevated β-cell fractional area in pancreases from Gα_z_ FΒKO females as compared to WT females (14). In our previous work, elevated β-cell fractional area correlated with elevated β-cell replication, as quantified by Ki67-positive, insulin-positive cells (14). Gα_z_ βKO, on the other hand, had no significant effect on β-cell replication in either sex (**Figure 5B**). When the percentage of Ki67-positive, insulin-positive cells is plotted on the same axis as in Fenske and colleagues to show the ~8% actively-replicating cells in Gα_z_ FΒKO female NOD mice(14), the lack of an effect on β-cell replication is apparent (**Figure 5C**).

**Figure 5:**
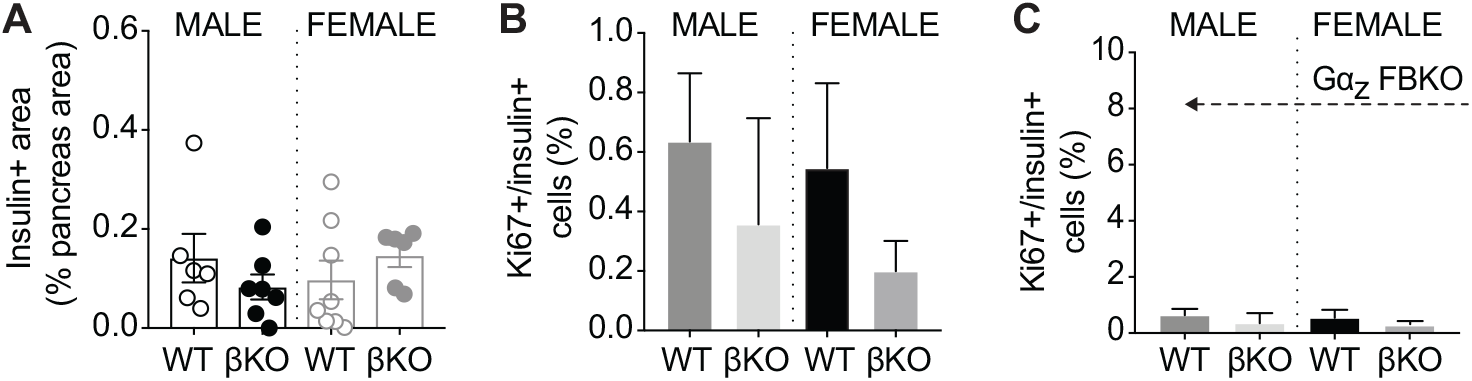
Loss of β-cell Gα_z_ does not impact overall β-cell area. A: Insulin-positive pancreas area of 30-35-week-old WT and Gα_z_ βKO mice as determined by insulin immunohistochemistry of pancreas sections. N=5-7. B: Percentage of Ki67-positive, insulin-positive cells in pancreas sections from 12-13-week-old male and female WT and βKO NOD mice. N=3-8. C: Same data as in (B) but plotted on the same y-axis as the results of Gα_z_ FBKO female mice as described in (14). The dashed arrow indicates the percent Ki67-positive, insulin-positive cells in same-aged Gα_z_ FBKO female NOD mice. In all panels, data represent means ± SEM. Data were compared by one-way ANOVA with Holm-Sidak test post-hoc to correct for multiple comparisons. No statistically significant differences were found.

### Loss of β-cell Gα_z_ does offer sufficient protection from a pure β-cell insult

Streptozotocin (STZ) is a recognized β-cell toxin that, when given using a low-dose protocol, induces β-cell death by apoptotic mechanisms!!ref. In the natural course of NOD diabetes development, β-cell survival is only a component of their relative susceptibility. In order to determine the importance of β-cell Gα_z_ in NOD β-cell apoptosis, we administered a low-dose (25 mg/kg) of STZ to 9-week-old NOD females for three consecutive days, instead of the typical 5 day protocol, to elicit a β-cell insult. Mice were followed for 5 weeks, until 14 weeks of age. As a control, the mean blood glucose levels of WT mice injected with citrate buffer remained near 100 mg/dl for the duration of the study, and no mice developed hyperglycemia during this timeframe (**Figure 6A,B**). The mean blood glucose levels of WT STZ-injected mice became elevated 1 week after STZ injection, and by 13-14 weeks of age, mean blood glucose was significantly elevated (**Figure 6A**). After 5 weeks, less than 25% of WT STZ-injected mice remained disease-free (**Figure 6B,D**). Both Gα_z_ βKO and Gα_z_ FΒKO mice maintained mean blood glucose levels at or below 250 mg/dl for the duration of the experiment (**Figure 6A,C**), and, compared to WT mice, had significantly improved and nearly-identical disease-free survival (**Figure 6D**).

**Figure 6:**
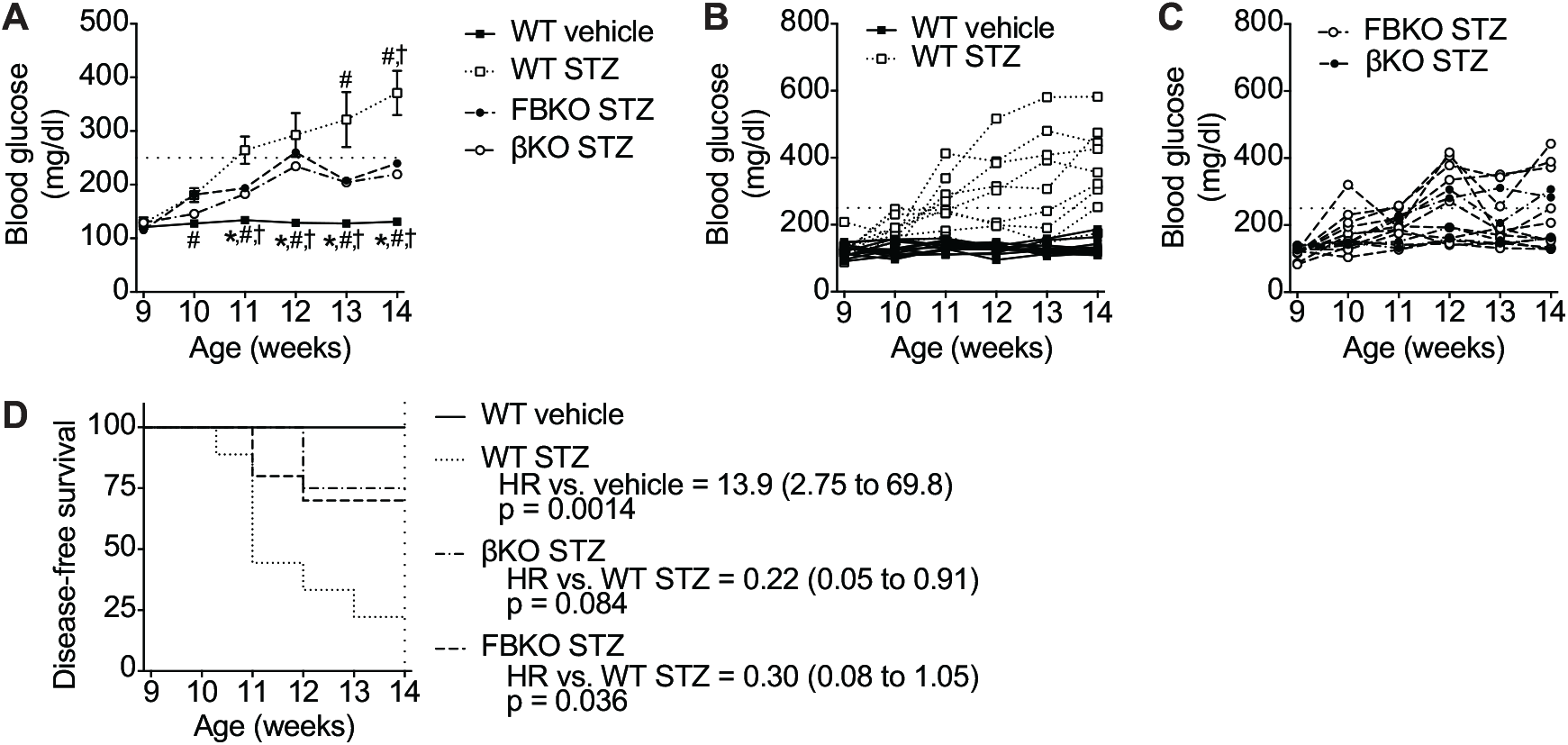
Beta-cell-specific Gα_z_ loss is sufficient to protect female NOD mice from a beta-cell insult. A: Mean Fasting blood glucose levels of WT, Gα_z_ FBKO, and Gα_z_ bKO mice after 3 consecutive days of injection with 50 mg/kg streptozotocin (STZ) or citrate buffer control (WT only). Data represent means ± SEM and were compared by two-way paired ANOVA with Holm-Sidak test post-hoc to correct for multiple comparisons. N=4-10. *p < 0.05 vs. WT STZ. ^#^, p<0.05 vs. Gα_z_ FBKO. ^†^, p<0.05 vs. Gα_z_ βKO. B,C: Weekly blood glucose levels of individual WT STZ and vehicle-control mice (B) and Gα_z_ FBKO and βKO mice (C). D: Disease free-survival of all four groups 5 weeks after STZ or vehicle administration. N=4-10.

## Discussion

Our lab has previously shown that global loss of Gα_z_ can fully protect female and male NOD mice from developing hyperglycemia (14). As the tissue distribution of Gα_z_ is extremely limited, we expected β-cell-specific Gα_z_ loss to phenocopy those of the FΒKO. Surprisingly, we found this was not the case. Loss of β-cell Gα_z_ was not sufficient to fully protect NOD mice from hyperglycemia and glucose intolerance. Furthermore, the effects of β-cell-specific Gα_z_ loss on different aspects of the NOD phenotype were sexually dimorphic, with βKO females having significantly improved glucose tolerance and βKO males having significantly reduced insulitis. Ultimately, the most striking difference between the results of the current work with our previously-published work is the lack of effect of β-cell-specific Gα_z_ loss on β-cell replication.

It has long been understood that male NOD mice have decreased prevalence of hyperglycemia than females, and, for that reason, many groups studying T1D in the NOD background use female animals only (23). The sex difference in phenotype penetrance in male and female NOD mice been attributed to a variety of factors, including prolactin levels, sex hormones, and gut microbiota (24–26). Our findings are confirmatory of those of previous works, with a 20% vs. 40% disease penetrance in male vs. female mice, respectively. Digging down into different aspects of the NOD phenotype, though, it does not follow that male NOD mice simply have a globally reduced magnitude of disease pathophysiology. In fact, and somewhat paradoxally, male NOD mice appear more glucose intolerant than females at 12-13 weeks of age, a timepoint that was chosen to precede the development of overt hyperglycemia in either sex. Furthermore, by 28-29 weeks of age, male WT mice have significantly better glucose tolerance than their younger cohorts, with higher plasma insulin levels as evidence of improved β-cell function. Female NOD mice, on the other hand, show little difference in *in vivo* metabolic phenotype by age, and it is only in the female sex that loss of β-cell Gα_z_ continues to provide a protective effect. The baseline insulitis score and the effect of β-cell Gα_z_ loss on islet insulitis also showed significant sexual dimorphism, and only in the male sex did insulitis correlate with glucose tolerance. Human T1D also presents in a sexually-dimorphic fashion, albeit with a higher prevalence in males than females (26). Therefore, even though the sex-dependence is reversed, performing full analyses in both NOD sexes could help to tease apart the sex-specific factors influencing T1D disease process, in mice and humans.

The effects of full-body vs. β-cell-specific loss of Gα_z_ have previously been characterized two different mouse models of diabetes: the high-fat diet (HFD)-induced model of T2D and MLD-STZ-induced model of T1D (13,14,16,19). While the protection of Gα_z_ βKO and FΒKO mice from HFD-induced T2D was nearly identical, the mechanisms behind this protection differed. Islets from HFD-fed Gα_z_ βKO C57BL/6J mice are significantly more incretin-responsive than those from WT HFD-fed mice, with no effect on β-cell replication or fractional area(19). This is in contrast to islets from HFD-fed Gα_z_ FΒKO C57BL/6N mice, in which the synergistic increase in β-cell replication by HFD feeding and Gα_z_ loss is the most striking aspect of the phenotype(16). In Schaid and colleagues, we proposed β-cell Gα_z_ may not directly influence β-cell replication at all, but may be permissive towards increased replication in the context of the relative insulin resistance of the C57BL/6N vs. C57BL/6J substrains(19). Like the HFD-induced model, the protective phenotype of Gα_z_ FΒKO C57BL/6N mice and Gα_z_ βKO C57BL/6J mice from MLD-STZ-induced T1D when mice are administered a sub-therapeutic dose of exendin-4 daily is nearly identical(13,14). Interestingly, though, in Brill and colleagues, where β-cell replication and apoptosis were both quantified, exendin-4 treatment and Gα_z_ loss elicited the same ~2-fold enhancement of β-cell Ki67 positivity, with no additional increase with the combination of the two. These results suggest a conservation of mechanism between the effect of islet Gα_z_ loss and GLP1-RA treatment on β-cell proliferation that is worthy of future exploration.

## Acknowledgements

We wish to thank the many present and former members of the Kimple Laboratory who contributed technical assistance or scientific discussion during the course of these experiments. This work was supported in part by Merit Review Award I01 BX003700 from the United States (U.S.) Department of Veterans Affairs Biomedical Laboratory Research and Development Service (BLR&D) (to MEK). Further support was provided by NIH Grants K01 DK080845 (to MEK), R01 DK102598 (to MEK), and F31 DK109698 (to RJF); ADA Grant 1-16-IBS-212 (to MEK), and JDRF Grant 17-2011-608 (to MEK). The content is solely the responsibility of the authors and does not necessarily represent the official views of the National Institutes of Health, the U.S. Department of Veterans Affairs, or the United States Government. The funding bodies had no role in any aspect of the work described in this manuscript.

## Author Contributions

Conceptualization, MEK; data curation, RJF, DCP, and MEK; formal analysis, RJF, DCP, HNW, and MEK; funding acquisition, RJF and MEK; investigation, RJF, DCP, HNW, MDS, AR, and KAK; methodology, RJF, DCP, HNW, KAC, and MEK; project administration, AMW, DBD, and MEK; supervision, NAT, RJF, AMW, DBD, and MEK; visualization, RJF, DCP, HNW, and MEK; writing—original draft, RJF and DCP; writing—review and editing, RJF, DCP, and MEK. All authors have read and agreed to the published version of the manuscript.

